# Individual Behavioral Variability Across Time and Contexts in *Dendrobates tinctorius* Poison Frogs

**DOI:** 10.1101/2023.09.25.559405

**Authors:** Katharina M Soto, Faith O Hardin, Harmen P Alleyne, Eva K Fischer

**Author notes:** Corresponding author: Eva K Fischer, Department of Evolution, Ecology, and Behavior School of Integrative Biology, University of Illinois Urbana-Champaign 505 S Goodwin Ave, Urbana, IL 61801.

## Abstract

Consistent individual differences in behavior (“animal personality”) have consequences for individual fitness, evolutionary trajectories, and species’ persistence. Such differences have been documented across a wide range of animals, though amphibians are generally underrepresented in this research area. The aim of our study was to examine consistent individual differences in Dyeing poison frogs, *Dendrobates tinctorius*. We evaluated repeatability in behaviors including activity, exploration, and boldness to assess consistency of behaviors across different temporal, experimental, and environmental contexts. We found repeatability in activity and exploration across time and contexts. In contrast, we observed context-specific behavior for our metrics of boldness, with consistent individual differences only for some measures. Further, while activity and exploration displayed consistent correlations across contexts, relationships between activity and boldness were context dependent. Our findings document the presence of consistent individual differences in behavior in *D. tinctorius* poison frogs, and also reveal context-dependent behavioral differences, highlighting the complex relationship between consistent individual differences and context-specific responses in animal behavior.

**Statement of Significance:** The concept of animal personality centers on the existence of consistent individual differences in behavior. However, behavioral responses can depend on context, and consistent individual differences in one context do not guarantee consistent differences in another. To address this question, we assessed activity, exploration, and boldness in captive-bred poison frogs (*Dendrobates tinctorius*) across time and environmental contexts. Our comprehensive approach revealed consistent individual differences in some behaviors and context-specificity in others. While activity and exploration were generally repeatable and correlated with one another, boldness was not. Especially in view of the emphasis on measures of boldness in the animal personality literature, our findings emphasize the importance of reiterative and holistic approaches in the study of animal behavior.

## Introduction

There is growing evidence that consistent individual differences in behavior exist across taxa, and that these differences can have important consequences for individual fitness, adaptation, and species’ persistence (Bell et al. 2009). For example, bold and aggressive individuals may be superior at territory defense but inferior at parental care (Duckworth 2006). Such trade-offs can lead to the establishment of alternative behavioral strategies within populations and potentially shape evolutionary trajectories (Smith and Blumstein 2008; Dochtermann and Dingemanse 2013). Moreover, certain personality types may fare better in the face of increasing human-induced global change, for example if they are better able to adapt to increasingly urban habitats (Pamela et al. 2015; Murgui and Hedblom 2017) or more likely to invade novel environments (Brodin et al. 2013; Carvalho et al. 2013). Increased recognition of the complex consequences of consistent individual behavioral differences have led to an increase in the study of animal personalities across taxa. Animal personality is increasingly invoked to explain behavioral differences in natural and captive populations, and to speculate on how these differences may influence survival and adaptation. However, many studies do so without confirming consistent individual differences across multiple contexts and over time.

Assuming – rather than testing – that individual behavioral variation is stable across time and contexts can introduce bias, leading to adaptive inferences based on isolated incidents or context-specific behavior (Bell et al. 2009). Moreover, design and interpretation of behavioral assays that facilitate cross-species behavioral comparisons remain a challenge (Uher 2008; O’Connell and Hofmann 2011; Odom et al. 2021).

Animal personality is broadly defined as consistent individual differences in behavior (Réale et al. 2010; Carere and Maestripieri 2013). These differences are typically characterized by quantifying behavioral repeatability, defined as the proportion of variance explained by within-individual variation compared to between-individual variation (Bell et al. 2009; Bergmüller 2010; Garamszegi and Herczeg 2012). Animal personality encompasses an array of behavioral traits, often with a particular focus on boldness:shyness, exploration:avoidance, and activity (Réale et al. 2007; Garamszegi and Herczeg 2012; Michelangeli et al. 2019). Further, these traits can become "packaged together” into behavioral syndromes (Sih et al. 2004). For instance, studies spanning diverse taxa such as common voles (*Microtus arvali*) (Herde and Eccard 2013), convict cichlids (*Amatitlania nigrofasciata*) (Mazué et al. 2015), sea turtles (*Chelonia mydas)* (Kudo et al. 2021), and great tits (*Parus major*) (Cole and Quinn 2011) document positive correlations between boldness, activity, and exploration.

While the presence of animal personalities and behavioral syndromes have been documented in animals from primates to beetles, there is bias in taxonomic representation. While there is extensive research in non-human animals including birds (e.g., Réale et al. 2007; Carere and Locurto 2011) and fish (e.g., Bell 2005; Smith and Blumstein 2010; Mazué et al. 2015), amphibians have received comparatively less attention. Indeed, recent cross-taxon reviews mention one or no studies on amphibians (Moiron et al. 2019; Cabrera et al. 2021). These biases likely arise in part from historical assumptions that amphibians were “largely instinctive bound machines controlled by external stimuli” (Maier and Schneirla 1964), yet it is increasingly clear that these creatures make complex, context dependent behavioral decisions (Burghardt 2013; Kelleher et al. 2018). Studying individual behavioral variation in amphibians is critical for advancing our understanding of how these animals navigate risks and opportunities, such as the trade-off between mating advertisement and predator avoidance (Stamps 2007; Sih and Del Giudice 2012). Importantly, this knowledge has broader implications for understanding evolutionary trajectories, ecological interactions, and conservation needs, which are of particular importance given the alarming global decline of amphibian populations (Kelleher et al. 2018).

Among anurans (frogs and toads), poison frogs (family Dendrobatidae) offer compelling study subjects for exploring individual variation and correlations in behavior. Poison frogs are known for their bright coloration, toxicity, and behavior. These species’ vivid hues are an honest signal of their toxicity and have coevolved with a diurnal lifestyle (Summers 2003; Maan and Cummings 2012). Poison frogs exhibit an array of interesting behaviors, including anti-predator defenses (Breed 2008), aggressive territoriality (Crothers and Cummings 2015), parental care (Roland and O’Connell 2015; Fischer et al. 2019), spatial learning and navigation (Liu et al. 2016; Pašukonis et al. 2022), and behavioral plasticity (Ringler et al. 2017; Peignier et al. 2023). Taken together, these characteristics have garnered poison frogs increasing attention in the realm of animal personality and behavioral syndromes. Existing studies in poison frogs have examined consistent individual differences in activity (Sonnleitner et al. 2022), antipredator behavior (Peignier et al. 2023), and aggression (Chaloupka et al. 2022). Nonetheless, a knowledge gap remains regarding the extent of consistent individual differences across time and context in poison frogs specifically, and amphibians more generally.

Dyeing poison frogs, *Dendrobates tinctorius*, have long been popular in zoos, aquaria, and the pet trade, and have recently received increasing research attention in the wild (Rojas and Pašukonis 2019; Fouilloux et al. 2021; Pašukonis et al. 2022) and the lab (Fischer et al. 2019, Fischer and O’Connell 2020; Sonnleitner et al. 2022). *D. tinctorius* boast a variety of bright color morphs without sexual dimorphism in coloration. In the wild, males and females maintain home ranges of a similar size, have similar daily movement patterns, and exhibit aggression toward same- and opposite-sex individuals (Rojas and Endler 2013; Pašukonis et al. 2022). Unusually among frogs, males of the species do not produce conspicuous advertisement calls, though they do produce low amplitude courtship vocalization (Rojas and Pašukonis 2019). Like other Dendrobatids, *D. tinctorius* provide extended parental care that includes defense, hydration, and cleaning of terrestrially deposited embryos, and transportation of tadpoles piggyback to pools of water upon hatching (Stynoski et al. 2015; Rojas and Pašukonis 2019). In *D. tinctorius*, tadpole transport is typically performed by males, however, females will occasionally take over (Rojas and Pašukonis 2019; Fischer and O’Connell 2020). Alongside their fascinating traits, existing work on *D. tinctorius* in both the field and the lab, and the ability to ethically acquire animals make them an excellent study system.

We investigated personality and behavioral syndromes across contexts and timeframes in *D. tinctorius*. Through a set of three experiments, we repeatedly assessed activity, exploration, and boldness – in the same and in different assays – to test whether commonly used personality metrics are robust to differences in assay design and timing (Fig. 1). Following the methodologies of Kelleher et al. (2018), our experiments included an open field test (OFT), a threat stimulus test (TST), a novel environment test (NET), and an activity in a familiar environment test (AFET) (Fig. 1). All experiments were built upon the open field test, a widely utilized method for assessing activity levels, exploratory behavior, and boldness across taxa (Bell et al. 2009; Carere and Locurto 2011). Importantly, individuals were assayed multiple times in each experiment (i.e., experienced multiple ‘rounds’ of each ‘trial type’ - terminology we use throughout the manuscript). This design allowed us to assess repeatability of commonly used behavioral metrics within and across contexts, and in response to multiple stimuli.

**Fig 1.**
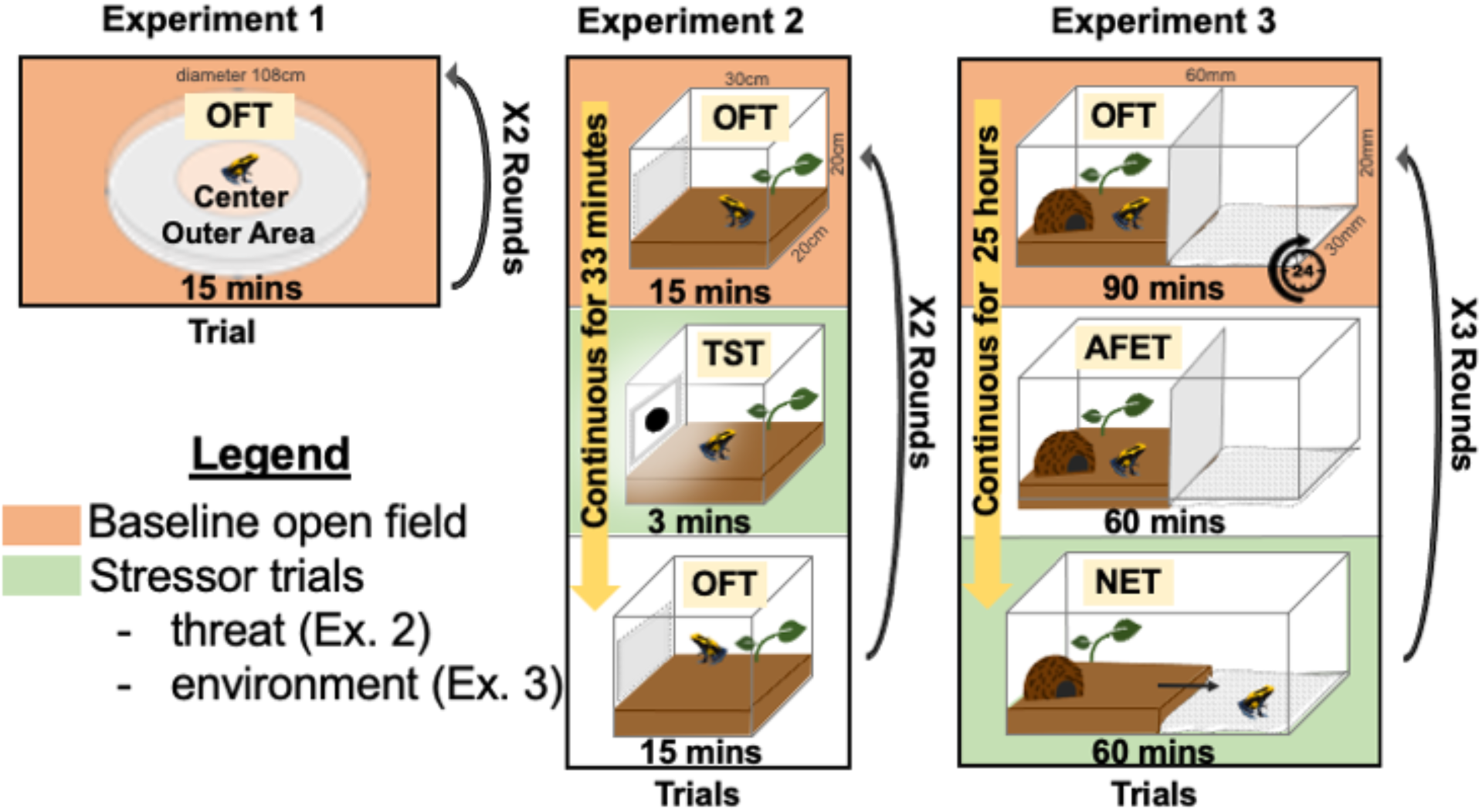
Overview of experimental design. All experiments were built on the framework of an open field test and expanded to explore differences in time and context. For each experiment, trial type (i.e., different phases within each experiment), trial duration, and number of rounds (i.e., repeated experiences in the same assay) are shown. OFT = open field experiment, TST = threat stimulus test, AFET = activity in a familiar environment test, NET = novel environment test. Colors represent corresponding trial types that were compared across experiments (see “Cross Analysis” Methods)

We predicted consistent individual differences in activity, exploration, and boldness across multiple rounds and the three distinct experiments. Further, we anticipated a behavioral syndrome, in which frogs with higher activity levels would also display increased boldness and exploration. Finally, we expected frogs to perceive the threat stimulus and the novel environment as stressful and exhibit context-dependent behavioral responses. Our overarching goal was to provide a comprehensive, nuanced understanding of how consistent individual differences in behavior manifest across time and context, and to reflect on the implications of these patterns for extrapolation to ecological and evolutionary outcomes.

## Methods

### Experimental overview

In Experiment 1, we conducted an open field test (OFT) in an artificial environment to establish baseline activity, exploration, boldness, and the relationships among them (Fig. 1). This test reflects the standard, basic design used for open field tests. In Experiment 2, we transitioned to an open field test (OFT) in a naturalistic environment, additionally assessing behavior before, during, and after exposure to a threat stimulus test (TST) (Fig. 1). In Experiment 3, we maintained the naturalistic environment, but extended the duration of each trial including the baseline open field test (OFT) and an activity in a familiar environment test (AFET), and additionally performed an extended duration novel environment test (NET) in an artificial environment (Fig. 1).

*D. tinctorius* were housed in our animal facility at the University of Illinois Urbana-Champaign. Terraria contained soil, moss, leaf litter, live plants, coconut shelters, and a small pond. For housing and all experiments, we maintained terraria and test arenas at 21°–23° C and 80%–90% humidity on a 12L:12D light cycle. We fed frogs *Drosophila hydei* or *Drosophila melanogaster* dusted with vitamins three times weekly.

All behavioral experiments were conducted between March and April 2022. We used a total of 48 captive bred frogs (N= 20 adult males, N= 12 adult females, and N= 17 juveniles of unknown sex). All frogs were assayed in Experiments 1 and 2. In Experiment 3, we used a subset of 19 frogs (N= 6 adult males, N= 8 adult females, and N= 5 juveniles; see additional details below). For Experiments 1 and 2, we ran frogs sequentially in groups of five per week, excluding the last group which had only four frogs. We followed the same protocol in a second round of trials with the same groups of frogs. In essence, each individual underwent two identical rounds, with a 5-week interval in between. For Experiment 3, we ran frogs in groups of nine per week through a series of sequential trials that lasted a combined 25 hours (details below). These trials were repeated three times with a 3-week interval between each round.

Group size for all experiments was chosen to make all behavioral testing feasible within a given daily time window and to balance individuals of different color morphs across groups. Group sizes and time intervals in Experiment 3 were modified to allow for the longer testing time course while still assaying the same frogs in all experiments within the same timeframe.

Although frogs varied in age, all individuals were sexually inexperienced. Previous studies have demonstrated that fed or digesting amphibians exhibit decreased levels of activity compared to fasted individuals (Andrade et al. 2005). To ensure that the frogs were not actively digesting during the experiments, we fasted frogs for 24 hours before behavioral trials.

### Experiment 1: Artificial open field test

In Experiment 1, we investigated exploration, activity, and boldness in an open field test (OFT) in a novel, artificial environment (Fig. 1). The open field arena was a circular enclosure with a diameter of 108 cm. To maintain humidity and moisture, we lined the arena with white bench paper soaked in reversed-osmosis (RO) water. For each observation, we placed a single frog at the center of the arena as a standardized starting point and recorded behavior for 15 minutes using an overhead camera (DVC, HDV-604S). Individuals were immediately returned to their home terraria following trials. The 15-minute testing time was chosen based on previous tests in amphibians and other species. We quantified exploration as the proportion of areas a frog visited, activity as the number of times a frog crossed the line between areas, and boldness as time spent in the center of area (Fig. 2) (additional details below).

**Fig 2.**
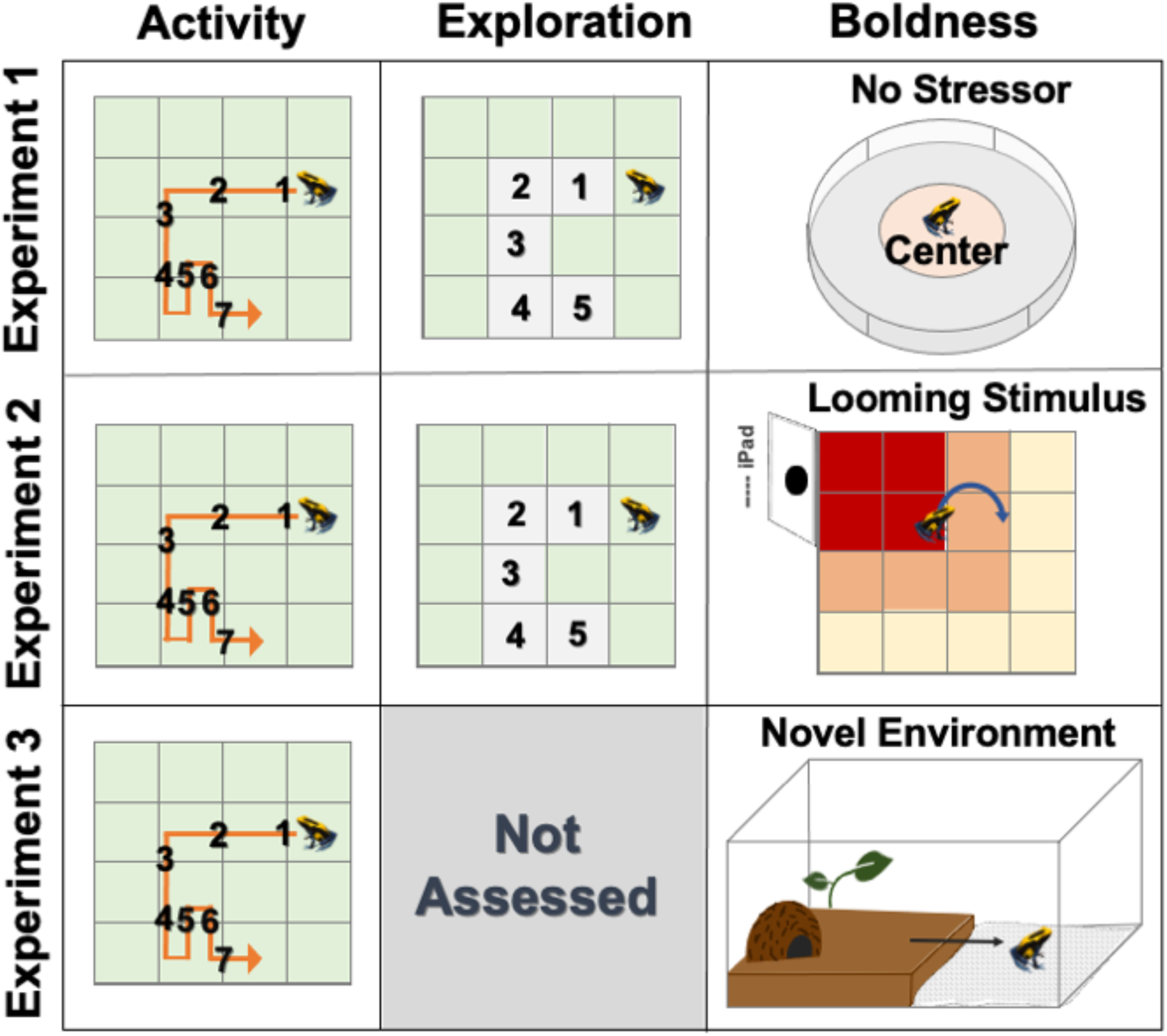
Overview of analysis methods. For all experiments, we quantified activity as the number of lines crossed from one area into the next and exploration as the total number of areas visited (not assessed in Experiment 3). Note that crossing back and forth from one area to another increases our measure of activity but not exploration. We quantified boldness as time in center (Experiment 1), response to a looming stimulus (Experiment 2), and latency to cross into a novel environment (Experiment 3)

### Experiment 2: Naturalistic open field and threat stimulus test

In Experiment 2, we quantified activity and exploration in a naturalistic open field. The behavioral arena was a 20 x 30 x 20 cm (L x W x H) plexiglass tank that mimicked the frogs’ home terraria, including soil substrate, moss, leaves, and a live plant positioned in one corner. The arena was completely covered from the outside with white contact paper, excluding a 24 x 17 cm (W x H) exposed area opposite the plant for the presentation of the threat stimulus. As in Experiment 1, we placed a single frog at the center of the arena at the beginning of each observation and used an overhead camera to record behavior. We conducted three trials to investigate behavior (1) pre-stimulus open field test (OFT) (15 minutes after introduction to arena), (2) during threat stimulus test (TST) (3 minutes of looming stimulus), and (3) post-stimulus open field test (for 15 minutes after the stimulus) (Fig.1). The 15-minute testing time was chosen based on previous studies and to align with Experiment 1. The duration of the looming stimulus was determined through pilot testing. Test trials on frogs not included in the study demonstrated that frogs typically showed a reaction within 3 minutes or froze throughout the entire stimulus presentation.

We assessed activity and exploration using the same methods as in Experiment 1 (Fig. 2). Additionally, we measured boldness by observing the frogs’ responses to a looming stimulus (Fig. 2). Previous studies using leopard frogs (*Lithobates pipiens*) have shown reliable avoidance behavior in response to looming objects (Ingle 1990; Waldeck and Gruberg 1995).

To present the looming stimulus, we used an iPad (Apple, 6th generation). The iPad was covered in white tape, except for the display screen, and placed in the exposed area of the arena (as shown Fig. 1). The looming stimulus was an expanding black circle projected repeatedly onto the iPad screen for a duration of three minutes. To prevent habituation and predictability, random pauses ranging from 1 to 30 seconds were incorporated into the stimulus presentation. During the pre- and post- exposure trials, the iPad displayed a white screen to match the surrounding environment. During the looming stimulus, we recorded activity and exploration, as well as boldness based on whether the frogs approached or displayed avoidance behavior in response to the stimulus.

### Experiment 3: Naturalistic open field, familiar environment, and novel environment tests

In Experiment 3, we increased the duration of each trial to capture more prolonged patterns in the frogs’ behavior. To conduct these longer trials concurrently with Experiments 1 and 2 while also maintaining our repeated measures design, we selected a subset of individuals (N= 19) from the previous experiments and limited our observations to two behaviors (activity and boldness). The behavioral arena was constructed using the same materials as Experiment 2. However, the dimensions of the arena were twice as long at 30 x 60 x 20 cm (L x W x H) and the arena split into two halves. One side of the arena mimicked the frogs’ home terraria as in Experiment 2. The other side mimicked the novel, artificial environment of Experiment 1, with only white, moistened paper flooring. The two sides of the arena were separated by a white cardboard barrier, which could be removed without disturbing the frog or the soil structure of the natural chamber.

We conducted three trials to investigate the frogs’ activity and boldness: (1) an open field test (OFT) in a novel, naturalistic environment (the first 90 minutes after introduction to the area), (2) activity in a familiar, natural environment test (AFET) (60 minutes after a 24 hour acclimation period in the arena), and (3) exposure to a novel, artificial environment test (NET) (60 minutes after the removal of the barrier between the familiar and novel environment) (Fig. 1). Time points were chosen to characterize differences based on longer observations (60 minutes = 4x the open field trial time in Experiments 1 and 2) and extended habituation to the test arena (and immediate 90-minute trial and a second trial after 24 hours). Trials were conducted sequentially, with each frog remaining in the arena for 25 hours while the trials progressed. We conducted two subsequent rounds of trials for each individual, with three weeks between each round. Due to the extended duration of each trial, we focused our analyses on activity and boldness. This decision was made to streamline data collection and because we expected – and found – a positive correlation between activity and exploration. We quantified activity using the same methods as in Experiment 1 and 2, number of line crosses. To assess boldness, we measured the frogs’ latency to cross into the novel environment when the barrier was lifted (Fig. 2).

### Data analysis

We describe general methods here with modifications for each experiment below. We coded videos using Behavioral Observation Research Interactive Software (BORIS) ver. 8.13 (Friard and Gamba 2016). Experiment type was not blind because assay identity was obvious from videos, but all analyses were blind to individual identity. Behavioral data was analyzed with R v.4.2.4 (R Core Team 2022) in Rstudio v.4.2.2. (RStudio Team 2021). For all response variables that were normally distributed, we used the raw data for analysis. Response variables that did not meet expectations of normality were transformed as described below. For all analyses, outcomes were considered significant at p <= 0.05 and all posthoc comparisons were Tukey adjusted for multiple hypothesis testing.

We used the lmer function in the lmerTest package (Kuznetsova et al. 2017) for all linear models, and always included frog ID as a random effect and round (i.e., repeated measurements in the same assay) as a fixed effect. Additional fixed effects varied based on experiment (see below). We tested for the effects of sex on behavior and found no significant results. Therefore, and because not all individuals could be sexed at the time of the experiment, we did not include sex as a factor in the analyses described below. We ran Type III ANOVAs on linear models to generate test statistics and p-values for main effects and interactions. Subsequently, we used the emmeans package (Lenth 2021) to run pairwise post-hoc comparisons where appropriate.

We performed repeatability analyses using the rptR package (Stoffel et al. 2017), with 95% confidence intervals (CI) based on 1000 bootstrapping runs. Frog ID was used as a random effect for repeatability (R) among individuals. We used Pearson correlation for normally distributed data and Kendall correlation for non-normally distributed data to assess relationships between behaviors. The correlation coefficients were interpreted using these criteria: ≈0.1 a small effect, ≈0.3 a moderate effect, and >0.5 a strong effect (Garamszegi et al. 2013).

### Analysis Experiment 1: Artificial open field test

We coded videos in BORIS using a transparent overlay that divided the behavioral arena into 40 areas: 24 on the ground and 16 on the wall. From videos, we quantified exploration as the proportion of areas a frog visited and activity as the number of times a frog crossed a line between areas. Movement between areas was recorded when the majority of the frogs’ body entered into a new area. We measured boldness as time spent in the center four areas of the arena (Fig. 2). We log transformed the activity and boldness data to improve normality. We ran linear models for each behavior (activity, exploration, boldness) as described above, including round as a fixed effect and frog ID as a random effect. We performed repeatability analysis with Gaussian distributions for exploration and Poisson distributions for activity and boldness (Stoffel et al. 2017). Data from 11 individuals was lost due to equipment failure, leaving a final sample size of N=38 individuals. Analyses and results are based on these samples.

### Analysis Experiment 2: Naturalistic open field threat stimulus test

We coded videos in BORIS using a transparent overlay that divided the behavioral arena into 32 areas: 16 on the ground and 16 on the wall. We quantified and analyzed activity and exploration as in Experiment 1, with the addition of a fixed effect of trial (pre-stimulus, during stimulus, and post-stimulus) to test for pre- and post- stimulus behavior differences. To compare trials of different duration, we divided each behavior by the total trial time.

For response to stimulus, which we used as a measure of boldness, we divided the arena into three sections: (1) at stimulus (the two wall areas overlaying the iPad, the two floor areas directly in front the iPad, and one wall area immediately to the left of the iPad), (2) near stimulus (the two wall areas and the four floor areas surrounding the “at stimulus” area), and (3) away from stimulus (the remaining areas) (Fig. 2). We noted the frogs’ position before and after stimulus presentation. We used a Chi-Squared test in the Tidyverse package (Wickham et al. 2019) for pairwise comparisons of responses (moved away, moved toward, or froze) (R Core Team 2022). We fit rptR models with Gaussian distributions to test for repeatability of each behavior across rounds and across trials (Stoffel et al. 2017).

### Analysis Experiment 3: Naturalistic open field novel environment test

We coded videos in BORIS using a transparent overlay that divided the behavioral arena into 96 areas: 16 areas on both the natural and novel arena floor, and 16 areas on the walls of each chamber. We recorded activity as the number of times a frog crossed the line between areas as in Experiments 1 and 2. After the divider was removed, we recorded the time it took for a frog to move into the novel area as latency to explore, which we interpreted as a measure of boldness (Fig. 2). We recorded activity on both sides of the chamber after the barrier was removed. We ran linear models for activity and latency to explore, including round (i.e., repeated measurements in the assay), trial (baseline, after acclimation, in a novel environment), and trial: round interaction as fixed effects, and frog ID as a random effect. As in Experiment 1, behaviors were divided by trial duration when analyzing trial effects.

To determine whether activity predicted an individual’s latency to cross into a new area, we filtered the data to only include observations taken during the novel environment test. We then fit a similar linear mixed effect model to predict latency to enter the novel area with fixed effects of activity, round, and their interaction, and frog ID as a random effect. We fit rptR models with Gaussian distributions to test for repeatability of each behavior across rounds and across trials (Stoffel et al. 2017).

### Cross-experiment analysis of activity and exploration

In addition to behavioral analyses within experiments, we were interested in the behavior of individuals across experiments. To compare exploration between Experiments 1 and 2, which had different arena sizes, we calculated the proportion of areas explored. We did not quantify exploration in Experiment 3, and it was therefore excluded from this analysis (Fig. 2). We compared the ratio of areas explored in Experiment 1 to both the pre-stimulus and post-stimulus exploration in Experiment 2 using linear mixed models, with exploration rate and experiment as fixed effects (Bates et al. 2015). We tested repeatability using a Gaussian distribution (Stoffel et al. 2017).

In all experiments, we measured activity by counting the number of line crosses (Fig. 2). To compare across experiments of different duration without introducing biases based on trial time, we divided the number of lines crossed by the total trial time to calculate a line crossing rate. We compared the subset of individuals (n= 19) in Experiment 3 with the same individuals from Experiments 1 and 2. Baseline activity measurements were similar across all experiments, taken prior to either the presentation of a stressor (threat stimulus test) or an environmental stressor (novel environment test) (Fig. 1). We also compared the post-stimulus activity in Experiment 2 to the activity observed during the novel environment test in Experiment 3, reasoning that these trials were similar in representing responses following an environmental change. We analyzed behavioral differences using activity rate as the response variable, experiment as a fixed effect, and frog ID as a random effect (Bates et al. 2015). We calculated post hoc comparisons and repeatability as before.

## Results

### Experiment 1: Artificial open field test

Frogs displayed repeatability in exploration (R= 0.278, p= 0.029) (Fig. 3a) and activity (R= 0.279, p= 0.032) (Fig. 3b) and across both rounds of an open field test separated by five weeks. However, time in the center of the arena, which we interpreted as a measure of boldness, was not repeatable (R= 0.044, p= 0.395). We found no significant effect of round for activity (F_1,37_= 3.390, p*=* 0.074), exploration (F_1,37_= 1.392, p*=* 0.245), or boldness (F_1,74_= 0.628, p*=* 0.431), suggesting that frogs’ behavior did not change in response to previous experience in the arena. Activity and exploration exhibited a strong positive correlation (corr= 0.767, p< 0.001). Boldness was not correlated with either activity (corr= -0.150, p= 0.196) or exploration (corr= -0.119, p= 0.307) (Table 1).

**Fig 3.**
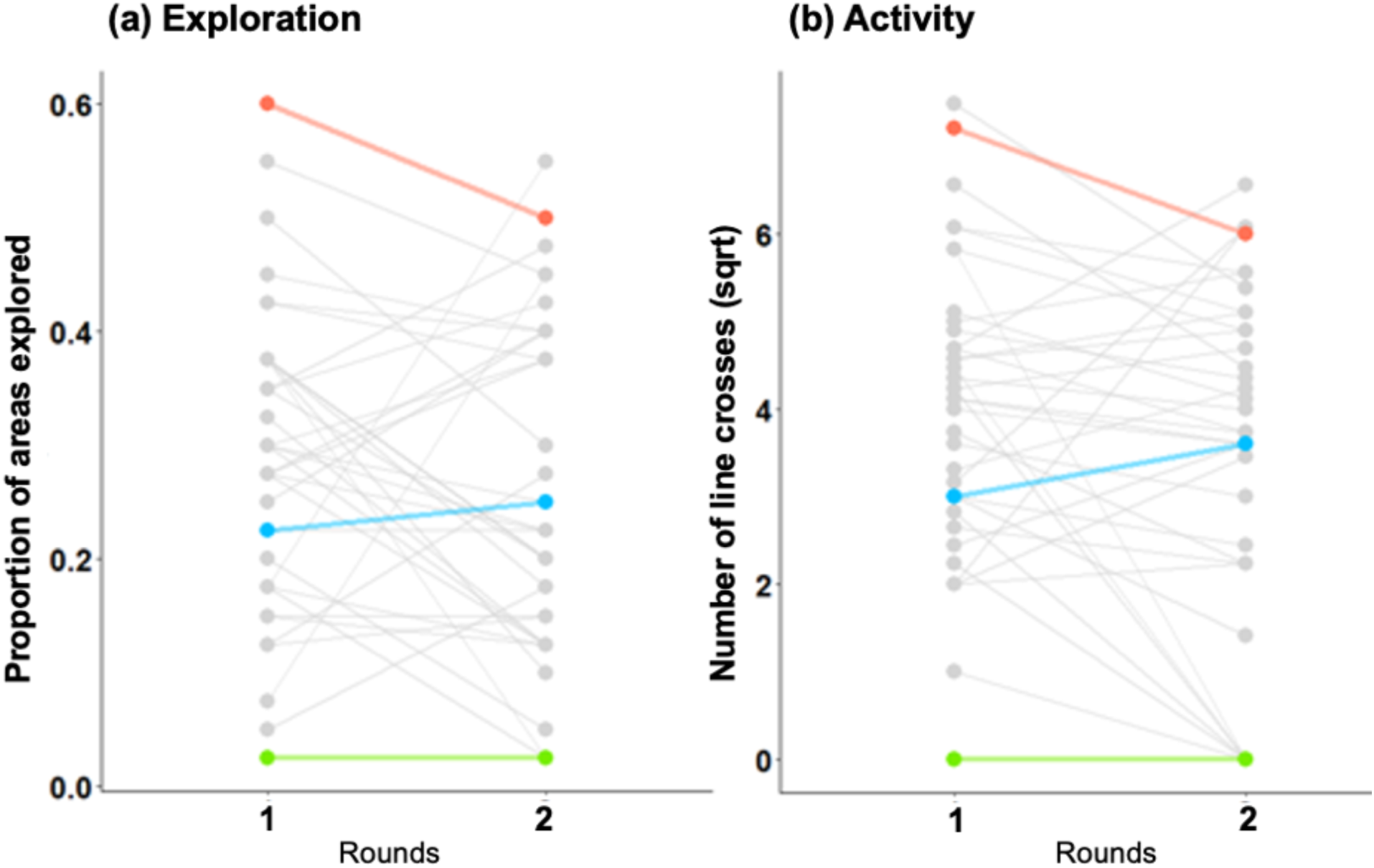
Behavior in an artificial environment open field test (Experiment 1). Frogs exhibited repeatability in (a) exploration (R= 0.278, p= 0.029) and (b) activity (R= 0.279, p= 0.032) across two rounds separated by five weeks. Three frogs are highlighted to showcase consistent individual differences in and the relationship between exploration and activity

**Table 1.**
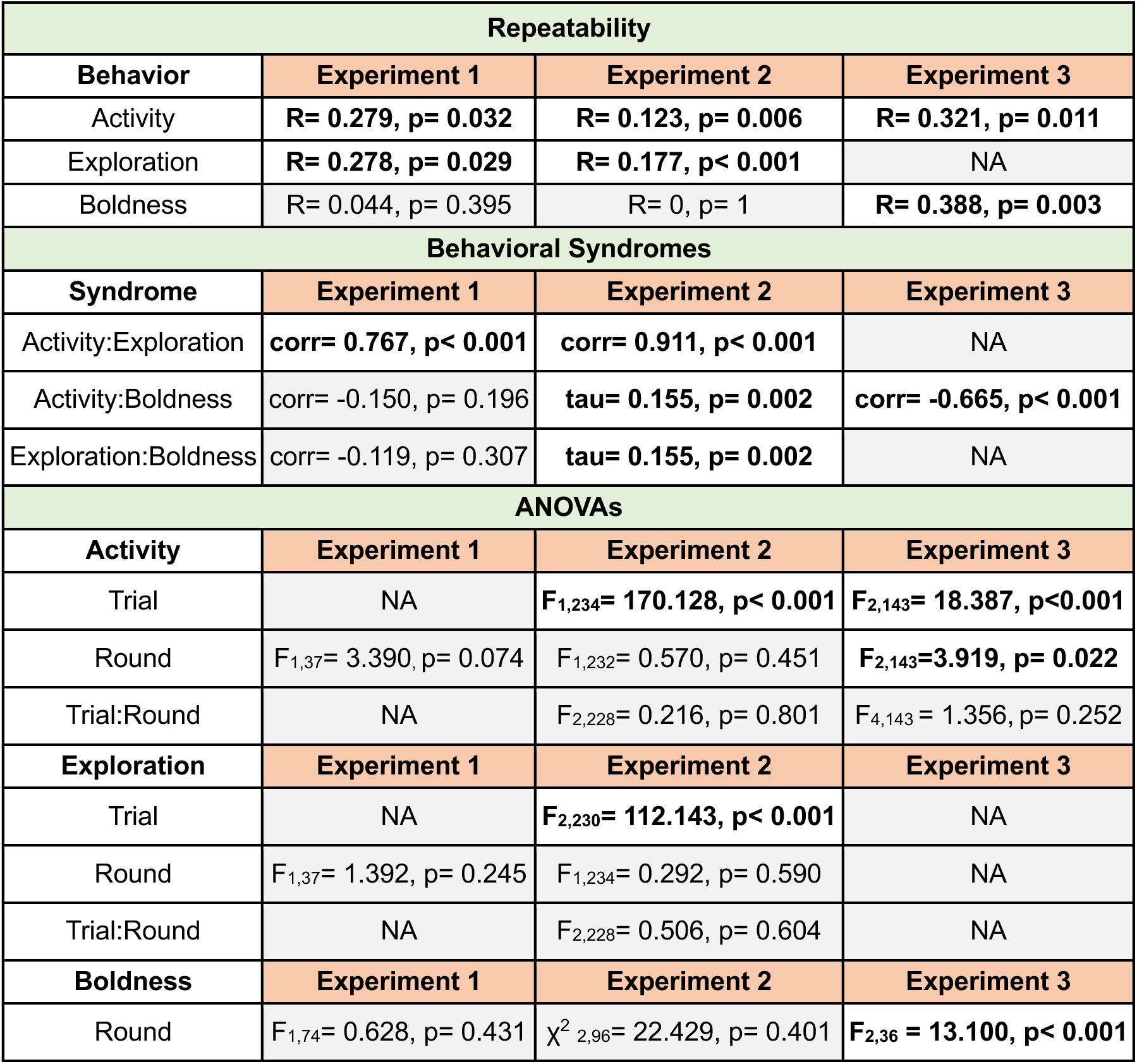
Overview of statistical results for behavioral repeatability, correlations, and linear models. Exploration was not quantified in Experiment 3. Significance interpreted as: R= 0.2-0.4 moderate repeatability, corr≥0.5 strong correlation, p< 0.05 significant. Pearson correlation (corr) used for normal data, Kendall Rank correlation (tau) used for non-parametric data

### Experiment 2: Naturalistic open field and threat stimulus test

Activity and exploration were repeatable across pre- and post- stimulus open field tests in both round 1 (activity: R= 0.593, p< 0.001; exploration: R= 0.421, p< 0.001) and round 2 (activity: R= 0.529, p< 0.001; exploration: R= 0.449, p< 0.001). We observed the same trend across rounds within trial (activity: R= 0.123, p= 0.006; exploration: R= 0.177, p< 0.001), aligning with the results from Experiment 1 (Table 1). In contrast, during the threat stimulus test, there was no repeatability of activity (R= 0.201, p= 0.069) or exploration (R= 0.175, p= 0.145) across rounds.

We found an effect of trial on exploration (F_2,230_= 112.143, p*<* 0.001) (Fig. 4a) and activity (F_1,234_= 170.128, p*<* 0.001) (Fig. 4b), with frogs decreasing behaviors during the threat stimulus test and then increasing post stimulus (Fig. 4). We observed no significant round effect for activity (F_1,232_= 0.570, p*=* 0.451) or exploration (F_1,234_= 0.292, p*=* 0.590), nor an interaction between round and trial (F_2,228_= 0.506, p= 0.604; F_2,228_= 0.216, p= 0.801), indicating that frogs behaved consistently across multiple experiences in the arena. Activity and exploration were highly correlated across all trials (pre-stimulus, during stimulus, and post-stimulus) (corr= 0.7– 0.8, p< 0.001).

**Fig 4.**
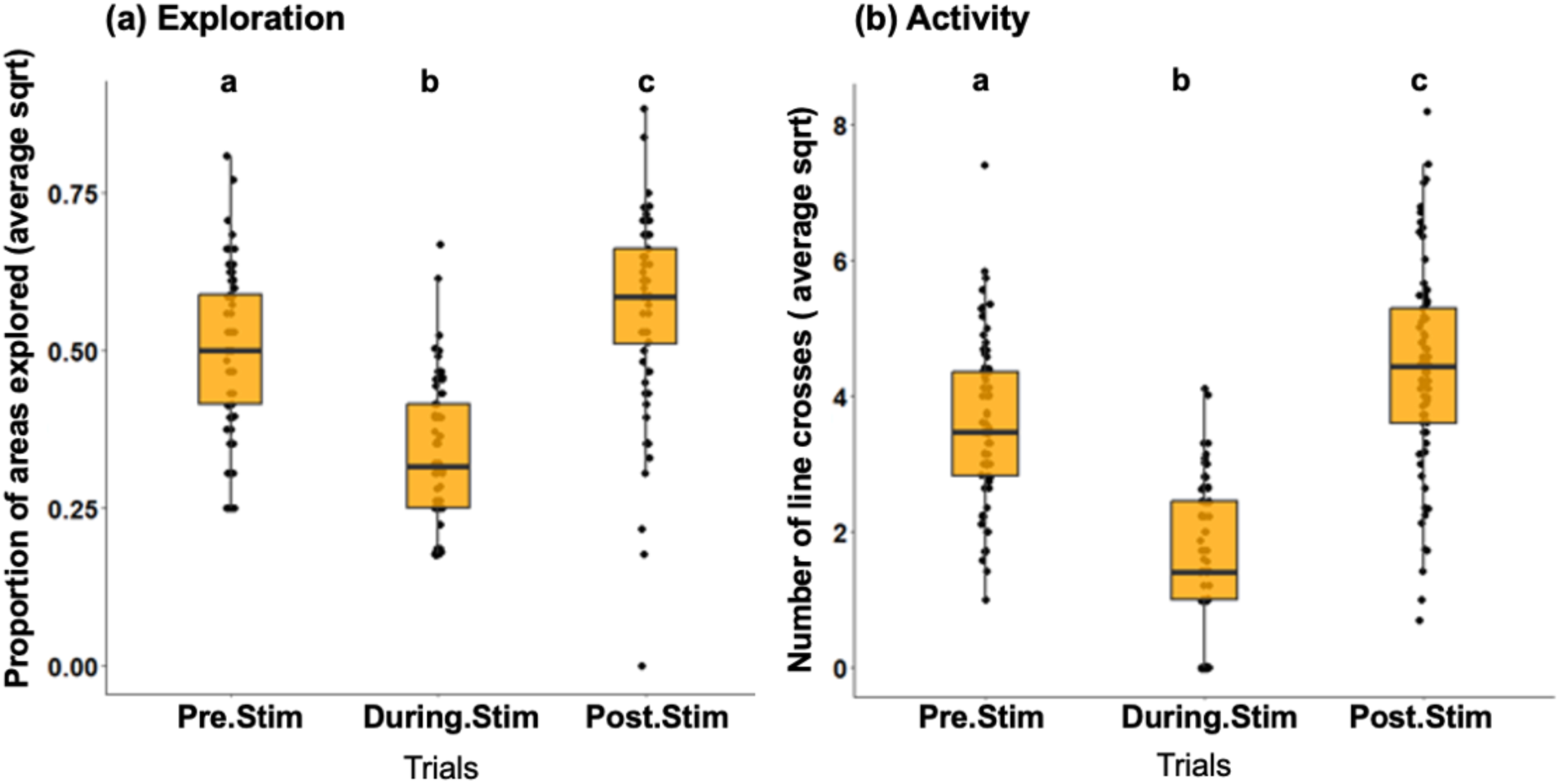
Behavior in a naturalistic open field before, during, and after a threat stimulus (Experiment 2). Exploration (a) and activity (b) decrease during stimulus presentation compared to pre- stimulus and post-stimulus (F_2,230_= 112.143, p*<* 0.001; F_1,234_= 170.128, p*<* 0.001), and these patterns were repeatable across the two rounds of the experiment. Data are collapsed across rounds, as no significant effect of round or a trial by round interaction were observed for exploration or activity. The boxes show the median, the first and third quartiles, and whiskers as 1.5x the interquartile range. Dots show individual data points.

We evaluated boldness by quantifying frogs’ responses to the threat (looming) stimulus. Frogs’ responses to the looming stimulus were not repeatable across rounds (R= 0, p= 1). This could be in part because overall among individual variation was low, as the majority of frogs demonstrated a freezing response during stimulus presentation rather than moving away (χ^2^_1,96_= 19.755, p*<* 0.001) or towards (χ^2^_1,96_= 41.796, p< 0.001) the stimulus. No significant effect of round was observed for boldness (χ^2^_2,96_= 22.429, p= 0.401).

We found significant correlations between boldness (response to stimulus) and frogs’ activity (tau= 0.155, p= 0.002) and exploration (tau= 0.155, p= 0.002) during the pre- and post-stimulus trials. These trends suggest that frogs displaying higher levels of activity and exploration prior to and after the stimulus were also more inclined to approach the stimulus during the threat stimulus trial. These correlations between boldness, activity, and exploration contrast with the results obtained in Experiment 1 (Table 1).

### Experiment 3: Naturalistic open field and novel environment test

Frogs were repeatable for activity both across trials (R= 0.321, p<0.011) and across rounds (R= 0.651, p= 0.001). We found that frogs moved more with repeated experience in the arena across all trials (F_2,143_= 3.919, p= 0.022), with round 1 having significantly less movement than round 2 (t_36_=-3.423, p=0.004) and Round 3 (t_36_= -3.784, p= 0.002), but no significant difference between rounds 2 and 3 (t_36_= -0.360 p=0.931) (Fig. 5a). Similarly, we found that frogs took less time to cross the boundary into the novel area during the novel environment test as rounds progressed (F_2,36_= 13.100, p< 0.001), with frogs taking longer to cross in round 1 compared to round 2 (t_36_= 2.837, p= 0.019) and round 3 (t_36_= 5.108, p< 0.001), but not in round 2 compared to round 3 (t_36_= 2.270, p= 0.073) (Fig 5b). We found significant repeatability for latency to explore across rounds (R= 0.388, p= 0.003) suggesting all frogs responded to repeated experience similarly.

**Fig 5.**
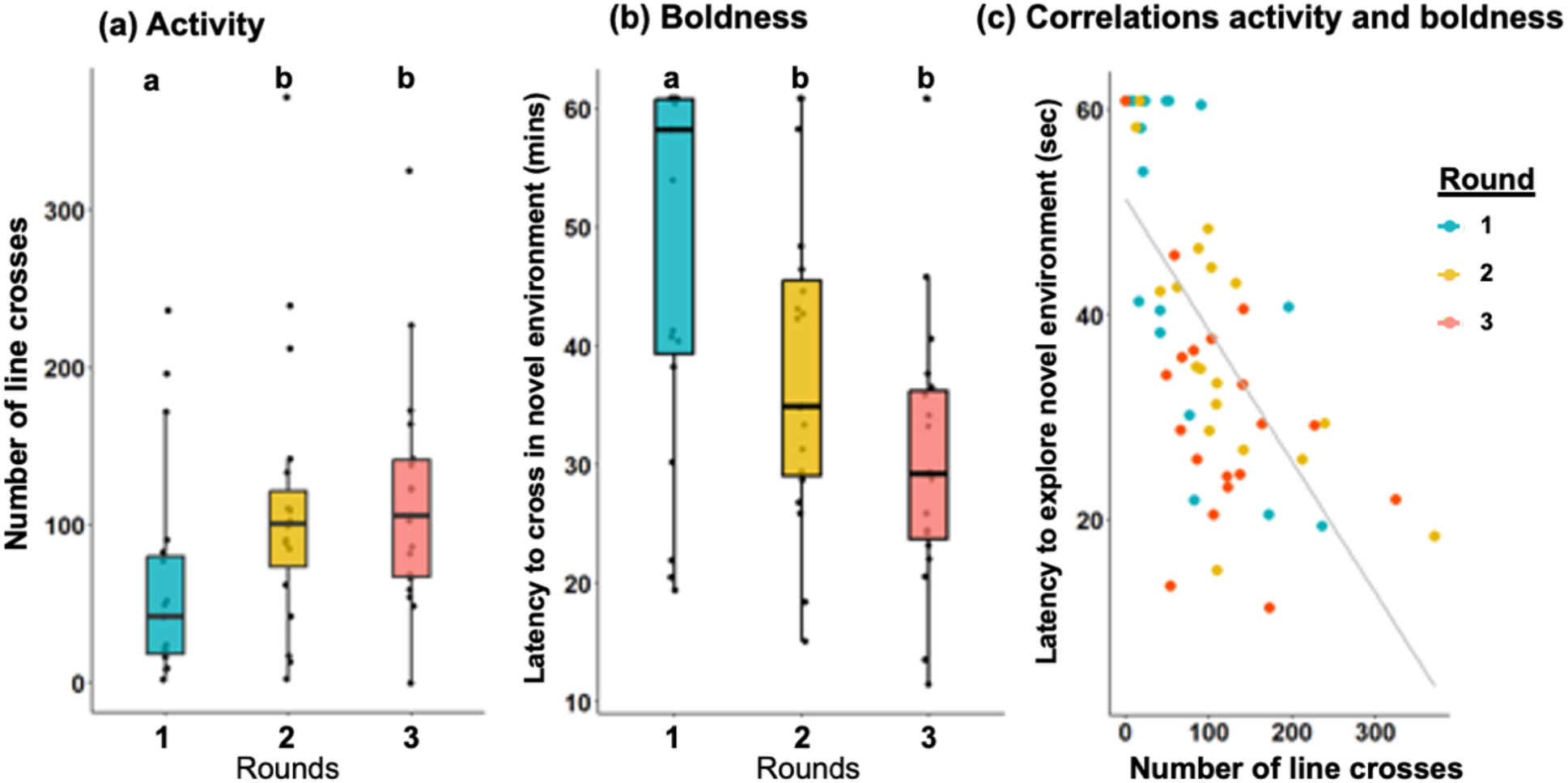
Behavior in a familiar, naturalistic environment and during the novel environment test (Experiment 3). (a) Activity increased with repeated experience in the assays (i.e., across rounds for the OFT, AFET, and NET; (F_2,143_= 3.919, p= 0.022). (b) All frogs explored the novel area more quickly in subsequent rounds (F_2,36_ = 13.100, p< 0.001), and (c) more active frogs explored the novel area sooner (corr = -0.655, p< 0.001). Letters in (a) and (b) indicate significant pair-wise differences between rounds. The boxes show the median, the first and third quartiles, and whiskers as 1.5x the interquartile range. Dots show individual data points.

We use latency to cross into a novel environment as an indicator of boldness, interpreting shorter latencies as bolder. We found a significant negative correlation between activity and latency to cross into a novel environment during the novel environment test (corr= - 0.665, p< 0.001), indicative of a positive relationship between activity and this measure of boldness (Fig 5c). Additionally, we found that boldness was influenced by both activity (F_1,38_= 28.795, p<0.001) and round (F_2,38_= 6.546, p= 0.004), indicating that more active frogs moved into the novel area more quickly, and all frogs moved into the novel area more quickly with repeated experience.

### Cross-experiment analysis of activity and exploration

Frogs exhibited consistent individual differences in exploration (R= 0.171, p= 0.005) across Experiments 1 and 2. For activity, baseline activity during the open field tests were repeatable between the naturalistic environments of Experiment 2 and Experiment 3 (R= 0.207, p< 0.001), and between the artificial environment of Experiment 1 and the naturalistic environment of Experiment 3 (R= 0.171, p= 0.041). Activity was not repeatable between the artificial environment of Experiment 1 and the naturalistic environment of Experiment 2 (R= 0.029, p= 0.401). Activity was repeatable across stressor trials (Fig. 1) (i.e., threat stimulus test in Experiment 2 and novel environment test in Experiment 3) (R= 0.243, p= 0.003).

## Discussion

We investigated animal personality and behavioral syndromes in captive-bred poison frogs (*Dendrobates tinctorius*) by characterizing behavioral differences across multiple environmental contexts and timepoints. We conducted three experiments to assess activity, exploration, and boldness in the same individuals and found repeatability in activity and exploration across time and contexts. In contrast, we observed context-specific behavior for our metrics of boldness, with consistent individual differences only for some measures. The stability of individual differences in behavior can have implications for how species respond to shifting environmental conditions (Wong and Candolin 2015; Merrick and Koprowski 2017), thereby shaping evolutionary trajectories and ecological interactions (Shaw 2020; Stiegler et al. 2022). Despite their declining populations, amphibians remain relatively understudied in behavioral research. The results of our work help to fill this gap, address broader issues in the field of “animal personality”, and set the stage for future work in *D. tinctorius*.

Our study revealed widespread repeatability for activity and exploration, evident across time, trials, and experimental contexts. These findings suggest that measures of activity and exploration are indeed reliable, and measurements in one assay or at one timepoint are good predictors of an individual’s behavior in another context or at another time. We also recovered the commonly described activity:exploration behavioral syndrome (Réale et al. 2007). While we expected a positive relationship between activity and exploration (i.e., exploration is impossible without movement), we note that there is room for variability in this relationship. For instance, frogs might have displayed a lot of movement but only within a restricted area, leading to heightened activity but limited exploration. The strong, significant relationships we found (correlations > 0.76), suggest that these behaviors are indeed tightly linked and can therefore reliably predict one another as well as some other behaviors (e.g., latency to explore a novel area, see below).

The above outcomes align with prior research on animal personality across taxa (Réale et al. 2007; Kelleher et al. 2018), and complement a recent investigation on another Dendrobatid, *Allobates femoralis*, which demonstrated the efficacy of both artificial and naturalistic setups in capturing consistent individual differences in behavior (Bégué et al. 2023). However, a study on *D. tinctorius* froglets by Sonnleitner et al. (2022) yielded somewhat different findings: the authors found significant repeatability at the family level but no significant repeatability at the individual level. These results suggests that genetic factors influence behavior, and this effect outweighs individual differences when related individuals are reared under similar conditions. We were unable to control for relatedness in our study, but assume genetic differences explain at least some of the behavioral variation we document. A non-mutually exclusive explanation is that individuals in the study of Sonnleitner et al. (2022) were much younger than those used here, and individual differences in behavior may become more pronounced with age and experience (Bell et al. 2009). We note that our findings here and those of Sonnleitner et al. (2022) are from captive-bred individuals. While *D. tinctorius* readily exhibit behaviors of interest (e.g., parental care and aggression) under captive conditions, influences of lab adaptation and genetic differences between wild and captive animals surely also influence behavior. In any case, while consistent individual differences are being increasingly documented across diverse taxa, contrasting results even in the same species underscore the importance of considering various contexts and timepoints when extrapolating from single studies to evolutionary and ecological outcomes.

While we observed a high degree of repeatability in activity and exploration across time and contexts, the behaviors we considered as measures of boldness demonstrated contextual dependence. Our first measure was the commonly used metric of time spent in the center of a novel arena (Carter et al. 2012; Yuen et al. 2017; Baker et al. 2018; Rudolfová et al. 2022). We did not observe consistent individual differences in this measure, nor was it correlated with other behaviors. As our next measure of boldness, we assayed response to a threat stimulus in a familiar environment, aligning with prior studies (Ingle 1990; Waldeck and Gruberg 1995; Carter et al. 2012; Yuen et al. 2017). Once again, we did not find repeatability for boldness, although we did observe a correlation between this measure of boldness, activity, and exploration. In our final approach, we measured boldness as the latency to enter a novel environment, a less common yet established method for evaluating boldness (White et al. 2013; Magnhagen et al. 2014; Theódórsson and Ólafsdóttir 2022). With this approach, we observed both repeatability and a correlation between boldness and activity. Taken together, our results suggest that commonly used boldness methodologies may not measure the same personality trait.

Despite the popularity of measuring boldness in animal personality studies, our findings align with those of others who have explicitly compared outcomes across different metrics categorized as ’boldness’ and found conflicting results. For example, our approach and outcomes are similar to that of Yuen et al. (2017) in their study on African striped mice (*Rhabdomys pumilio*). The authors investigated whether distinct experimental methods were indeed measuring boldness. Like our approach, they employed both artificial and natural environments, and conducted two boldness tests, an open field, and a startle test. Yuen et al. (2017) highlighted the distinction between these methods: the startle test evaluated risk-taking behavior after threat exposure, potentially measuring anxiety rather than boldness, while the open field test did not incorporate a preceding threat and proved to be a more reliable method for gauging boldness in their species. Carter et al. (2012) found related results when examining chacma baboons (*Papio ursinus*), concluding that threatening situations may measure anxiety, not boldness. Moreover, the use of artificial open field tests, particularly in evaluation of time spent in the center, may not consistently capture boldness. Indeed, a recent study on Zebrafish (*Danio rerio*) suggests that thigmotaxis may not effectively measure boldness or anxiety-like behavior in this species (Rajput et al. 2022).

Translating these observations to our own results, our open field test focused on evaluating time in center, the threat stimulus test observed behavior in the presence of a threat, whereas the novel environment test examined behavior in a new and unfamiliar environment.

While we initially considered all tests as measures of boldness, our data suggest that the specific nature of the challenges presented in each experiment and the environmental context led to varying types and intensities of behavioral responses, and in turn distinct patterns of repeatability and behavioral correlations. Our findings also confirm the importance of considering the environment when measuring behavioral correlations: we observed correlations between boldness, activity, and exploration (i.e., behavioral syndromes) in naturalistic environments (Experiments 2 and 3) but not in artificial settings (Experiment 1) (Table 1).

Alongside those of others, our findings highlight that the environment and assay in which individuals are assessed can exert a profound influence on their behavior (e.g., Niemelä and Dingemanse 2014; Bégué et al. 2023) and the necessity of evaluating individual variation across diverse contexts (Bell et al. 2009).

Taken together, these observations emphasize the challenges of adapting standard methodologies across closely and distantly related taxa. For example, poison frogs are diurnal unlike most frogs, and amphibians are generally less active than mammals. While behavioral and life-history differences between taxa are obvious and themselves of interest, they provide fundamental challenges for the design, implementation, and interpretation of comparative studies (Uher 2008; O’Connell and Hofmann 2011; Odom et al. 2021). When findings align across taxa, we assume shared core principles; however, when patterns differ it is more difficult to understand whether organisms differ fundamentally in, for example, relationships between activity and boldness, or whether we have not “asked them the right question” with our chosen assay. Fortunately, with a growing literature that includes more species and taxa, these challenges are also opportunities to build our understanding through comparative data and thinking.

## Conclusions

Our findings demonstrate consistent individual differences in behavior across time and context in *D. tinctorius*, while also highlighting the subtle complexities of studying animal personality. While we found strong evidence for individual consistency in activity and exploration, our measures of boldness were variable and context dependent. Our results demonstrate the value of assessing multiple behavioral traits and the relationships among them to build a comprehensive view of behavioral repeatability and variability within and across species (Sih et al. 2004; Bell et al. 2009; Sih and Del Giudice 2012). Broadly, our results add to the growing literature emphasizing the importance of evaluating context-dependent patterns of individual behavioral variation, even for assays commonly considered as measuring the same trait (Bell et al. 2009; Carter et al. 2012; Yuen et al. 2017). More specifically, our findings contribute to the growing recognition of behavioral complexity in anuran amphibians, with implications for amphibian research, including their role as ecological indicators amid habitat degradation and climate change (Grant et al. 2019). Our findings here pave the way for future research on the impact of consistent individual differences on reproductive success, survival, and community dynamics.

## Acknowledgments

We thank the members of the Fischer Lab for feedback on previous versions of the manuscript and help with frog husbandry. For their insight on statistical analyses, we thank Drs. Jen Moss and Sarah Westrick. We thank two anonymous reviewers and the associate editor for comments that improved the manuscript. Our work would not be possible without the animal care support of the Division of Animal Resources (DAR) at UIUC.

## Data availability

Data are included in the Supplementary information.

## Author Contributions

KMS, FOH, HPA, and EKF conceived of and planned the experiments. KMS, FOH, and HPA carried out the experiments. KMS, FOH, and EKF contributed to data analysis and interpretation of the results. KMS wrote the manuscript with assistance from all authors.

## Compliance with Ethical Standards

### Ethics approval

All animal care and experimental procedures were approved by the University of Illinois Urbana- Champaign Animal Care and Use Committee (protocol #20147) and followed all institutional and national guidelines for the use of animals in research.

